# Features of the DNA *Escherichia coli* RecN interaction revealed by fluorescence microscopy and single-molecule methods

**DOI:** 10.1101/2024.04.09.588753

**Authors:** Viktoria D. Roshektaeva, Aleksandr A. Alekseev, Alexey D. Vedyaykin, Viktor A. Vinnik, Dmitrii M. Baitin, Irina V. Bakhlanova, Georgii E. Pobegalov, Mikhail A. Khodorkovskii, Natalia E. Morozova

## Abstract

The SOS response is a condition that occurs in bacterial cells after DNA damage. In this state, the bacterium is able to recover the integrity of its genome. Due to the increased level of mutagenesis in cells during the repair of DNA double-strand breaks, the SOS response is also an important mechanism for bacterial adaptation to the antibiotics. One of the key proteins of the SOS response is the SMC-like protein RecN, which helps the RecA recombinase to find a homologous DNA template for repair. In this work, the localization of the recombinant RecN protein in living *Escherichia coli* cells was revealed using fluorescence microscopy. It has been shown that the RecN, outside the SOS response, is predominantly localized at the poles of the cell, and in dividing cells, also localized at the center. Using *in vitro* methods including fluorescence microscopy and optical tweezers, we show that RecN predominantly binds single-stranded DNA in an ATP-dependent manner. RecN has both intrinsic and single-stranded DNA-stimulated ATPase activity. The results of this work may be useful for better understanding of the SOS response mechanism and homologous recombination process.

## 1. Introduction

DNA recombination systems are important for living organisms to maintain the integrity of their genome. The genome integrity is critical for the life of all organisms, while DNA damage occurs regularly due to various factors, such as exposure to endogenous and exogenous damaging agents [1], ionizing radiation, UV radiation and others [2]. Double-strand DNA breaks can be lethal if they are not corrected before cell division, so both bacteria and eukaryotes have several mechanisms to repair such damage. In bacteria, depending on the microorganism, the repair of double-stranded DNA breaks can be carried out by the RecBCD, AddAB and AdnAB complexes. These multisubunit complexes form unique helicase-nuclease machines that process damaged DNA ends and load the RecA protein at the break site, which then initiates homologous recombination [3].

The RecA protein is a recombinase that is a key participant in homologous recombination DNA repair in bacteria. RecA activity has complex, multilevel regulation. Expression of the RecA gene in bacteria increases at the very first stage of a special cell state caused by DNA damage, called the SOS response [4]. Another protein that is highly induced at the early stage of the SOS-response is RecN, an SMC-like protein that facilitates repair of double-strand DNA breaks [5]. Upon DNA damage, RecN forms foci on DNA and colocalize with RecA, apparently representing repair centers [6].

Live cell imaging of *C. crescentus* revealed that RecA filaments exhibit directional translocation during homology search, which depends on RecN and its ATPase activity. *In vitro* studies of purified *D. radiodurans* proteins also showed that RecN facilitates the ligation of linear duplex DNA molecules [7] and stimulates RecA-mediated DNA strand exchange [8]. In addition, RecA-DNA complex stimulates ATPase activity of RecN, significantly changing the kinetics of the ATP hydrolysis [8]. RecN has also been shown to be a DNA-binding protein that preferentially binds ssDNA [9]. Interestingly, RecN apparently is required for the reorganization and compaction of the nucleoid after DNA damage [10].

RecN is one of the key factors of the SOS response and an important mediator of the homology search during homologous recombination in bacteria. However, the exact role of RecN in RecA-mediated double-strand break repair is not fully understood. Difficulties in obtaining a soluble recombinant *E. coli* RecN obstructed its detailed biochemical characterization. Recently an active recombinant *E. coli* RecN was obtained for the first time [9], which opens new opportunities for further research into how RecN functions *in vitro*. This article develops the topic and explores RecN-DNA interactions and the role of its ATPase activity.

## 2. Materials and methods

### 2.1 Strains, plasmids and growth conditions

Two genetic constructs were created. First *recN* gene was amplified from *E. coli* MG1655 strain. At the next step *E. coli recN, recN* fused to mCherry and only mCherry gene were placed into standard cloning vector pET21a under control of Lac operon. The fusion protein was made by adding mCherry to the C-terminus of RecN. The cloning procedure was made using restriction-ligation technique by XhoI and BamHI restriction sites. The design of genetic constructs assumes a polyhistidine tag at the C terminus of RecN or RecN::mCherry which allows proteins purification using Ni-NTA affinity chromatography. All cloning procedures were performed using *E. coli* DH5λ strain. Since expression of *recN* is under control of T7 promoter, RecN, mCherry and RecN::mCherry proteins production were performed into *E. coli* Rosetta (DE3) strain.

Growth conditions for RecN and RecN::mCherry proteins production for subsequent purification were as follows: an overnight culture of *E. coli* Rosetta (DE3) cells carrying pET21a_RecN or pET21a_RecN::mCherry plasmids were diluted 1:100 in 500 ml of fresh LB medium containing selective marker ampicillin. The fresh cell culture was grown at 37°C until OD_600_ reached 0.6. Expression of RecN or RecN::mCherry was then induced by 1 mM IPTG, then the cells were grown for 2 hours. Further, cell cultures were pelleted by centrifugation for 15 minutes at 4000 g at 4°C. Palettes then were used for RecN and RecN::mCherry purification.

To study localization of RecN inside the individual cells out of SOS response, fluorescence microscopy of living *E. coli* Rosetta carrying pET21a_RecN::mCherry and pET21a_mCherry plasmids was performed. To visualize RecN localization an overnight culture of *E. coli* Rosetta (DE3) carrying pET21a_RecN::mCherry or pET21a_mCherry plasmid was added 1:100 to fresh LB medium containing selective marker ampicillin. The fresh cell culture was grown at 37°C until OD_600_ reached 0.3 and then RecN::mCherry or mCherry expression was induced with 1mM IPTG. Next cells were grown for additional 1 hour at 37°C before imaging.

### 2.2 Recombination activity of the RecN

The recombination activity of RecN was evaluated. The recombination activity of the RecN protein in *E. coli* were determined in conjugation experiments, followed by calculation of the yield of recombinants and the frequency of recombination exchanges [11].

### 2.3 Purification of RecN and RecN::mCherry proteins

To purify the RecN and RecN::mCherry proteins, we used the buffer system described in [9]. However, to optimize existing buffer system, Buffer B (20mM Tris–HCl at pH 7.5, 1M NaCl, 1 mM EDTA, 10% glycerol, 7 mM 2-mercaptoethanol, 1 mM phenylmethylsulfonyl fluoride, and 0.3mM Triton X-100) from [9] was supplemented with 0.3mM of Triton X-100.

Briefly, cell pellets were dissolved in 20 ml of modified buffer B (20mM Tris–HCl at pH 7.5, 1M NaCl, 1 mM EDTA, 10% glycerol, 7 mM 2-mercaptoethanol, 1 mM phenylmethylsulfonyl fluoride, and 0.3mM Triton X-100). The cell suspensions were lyzed with ultrasound until the lysate was clarified. Next, the suspension was centrifuged at 4°C (12000g, 10 minutes), the supernatant was collected and passed through a syringe filter with a pore size of 0.45 μm. The proteins were purified using an Akta Purifier 10 chromatograph (GE Healthcare) and His Trap column. Proteins were eluted from the column using a linear gradient of 70–500 mM imidazole. All samples were applied to a polyacrylamide gel for subsequent protein electrophoresis and determination of fractions containing the largest amount of protein. Selected fractions were concentrated in centricons (Amicon).

The RecN and RecN::mCherry concentrations were estimated spectrophotometrically by absorption at a wavelength of 280 nm, adjusted for molecular weight and the molar extinction coefficient of the RecN and RecN::mCherry proteins: MW=63 kDa, 25900 cm-1•M-1 and MW= 90 kDa, 60280 cm-1•M-1, respectively.

### 2.4 Visualization of RecN::mCherry localization in living bacterial cells using fluorescence microscopy

In microscopy experiments of cells expressing RecN::mCherry fusion DNA of bacteria was stained with the intercalating dye DAPI (Thermo). Stained bacteria were placed on microscope slides preparation of which was described previously [12].

Fluorescence microscopy was performed using a Nikon Eclipse Ti-E inverted microscope under control of Micro-Manager 2.0 software. Images were collected in three channels: in transmitted light mode, in the red fluorescent channel (Semrock TxRed-4040C) and in the violet fluorescent channel (Semrock DAPI-50LP-A). All images were acquired using a Zyla 4.2 sCMOS camera (Andor) and were analyzed using ImageJ software.

### 2.5 Determination of *E. coli* RecN ATPase activity

The ATPase activity of the RecN protein was determined using a spectrophotometric method (Varian Cary 5000) [13]. Measurements were carried out at 37°C in 200 μl of a reaction mixture with 20 mM Tris-OAc (pH 7.5), 10 mM Mg(OAc)2, 80 mM KOAc, 2 mM ATP, 2 mM phosphoenolpyruvate, 30 U/ml pyruvate kinase, 30 U/ml lactate dehydrogenase, ∼0.2 mM NADH and 3 μM RecN. To determine the effect of DNA on the RecN ATPase activity, 6 μM poly(dT) (concentration relative to the number of bases) was also added to the reaction. The formation of ADP during ATP hydrolysis was chemically coupled to the oxidation of NADH to NAD+ and was recorded by a decrease in absorption at a wavelength of 340 nm using the NADH extinction coefficient ε340 = 6.22 mM-1•cm-1.

### 2.6 Visualization of RecN::mCherry interactions with single DNA molecules straightened on a substrate

To visualize binding activity of RecN::mCherry to various substrates, the protein (10^3^ protein molecules per 1 DNA molecule) was incubated with ssDNA or dsDNA (about 9600 bp length) at 37°C for 10 minutes. The incubation was performed in a buffer solution composed of 20 mM Tris-HCl, 7.7 mM KCl, 2.5 mM MgCl2, 1 mM DTT (pH=7.0), 2 mM ATP. After incubation with protein, DNA was stained with the green DNA-binding dye YOYO-1 at a concentration of 10 nM.

Cover slips (22x22 mm, Menzel-Gläser) were coated with polystyrene using a spin coater. Then, using the molecular combing technique [14], DNA-RecN mixture was straightened on a glass surface. Glass slides (Menzel-Gläser) were used to form microscope slides. A strip of double-sided adhesive tape (Scotch) was glued onto the slide to serve as a separator between the slide and a cover slip. A cover glass with straightened DNA was attached to adhesive tape, and a chamber was formed between the two glasses into which the buffer solution was washed. Fluorescence images were recorded as described in 2.4. The green fluorescence was detected using Semrock YFP-2427B filter set.

### 2.7 Visualization of RecN::mCherry DNA interaction dynamics at the level of individual molecules

Optical trapping experiments were performed to visualize RecN::mCherry protein binding to individual double-stranded and single-stranded DNA molecules. DNA manipulation was carried out inside a five-channel microfluidic system (uFlux, Lumicks), using two polystyrene microspheres coated with streptavidin (Spherotech, 2.1 μm diameter) and held by two optical traps according to the method presented in [15,16]. Double-stranded linear DNA molecules ∼23 kb length with biotinylated ends and obtained by modifying the DNA of bacteriophage lambda were initially used as a DNA substrate. The DNA molecule was attached to the microspheres through biotin-streptavidin interaction. Single-stranded DNA molecules were obtained directly during the experiment from double-stranded DNA molecules by force melting.

The force melting method is based on stretching a double-stranded DNA molecule with a force of more than 80 pN, resulting in the breaking of hydrogen bonds between complementary strands. Due to the modification of the 5’- and 3’-ends of the same strand of a double-stranded molecule by biotin, DNA melting leads to the fact that only one strand of DNA remains between the microspheres; the strand complementary to it is irreversibly removed from the held microspheres due to diffusion.

To visualize RecN::mCherry binding to individual DNA molecules, the following solutions were placed into the channels of the microfluidic chamber: 1st channel: 0.01% solution of polystyrene streptavidin-coated microspheres; 2nd channel: 3.5 pM solution of double-stranded DNA; 3rd channel: buffer solution; 4th channel: buffer solution with 1 mM ATP; 5th channel: 400 nM RecN::mCherry, 3 mM ATP, 10 U/ml pyruvate kinase, 1 mM phosphoenolpyruvate.

The buffer solution in all channels was 25 mM Tris-HCl (pH 7.5), 10 mM NaCl, 5 mM MgCl2. The experiments were carried out at a temperature of 22°C. The capture of microspheres, the attachment of a single DNA molecule to them, and the conversion of DNA into a single-stranded form by force melting were carried out in the first three channels of the microfluidic chamber. Individual single-stranded and double-stranded DNA molecules were incubated in a channel with 400 nM RecN::mCherry for 3 minutes, after which they were transferred into the channel of a microfluidic chamber containing a buffer solution without RecN::mCherry and fluorescent visualization of DNA-protein complexes was carried out.

Fluorescence was excited using a diode-pumped solid-state laser with a wavelength of 532 nm. Fluorescence was separated from the exciting radiation using a Filter set 10 from Carl Zeiss. Images were collected using EMCCD camera (Cascade II, Photometrics).

### 2.8 Analysis of DNA electrophoretic mobility shift in the presence of RecN

To confirm that *E. coli* RecN binds DNA, an electrophoretic mobility assay of DNA in the presence of the protein was performed on an agarose gel. Plasmid DNA (∼9600 bp) was used as a substrate into two forms: circular single-stranded and circular double-stranded DNA. Buffer containing 20 mM Tris-HCl (pH=7), 7.7 mM KCl, 2.5 mM MgCl2, 1 mM DTT, was used. Single-stranded DNA was prepared by heating at 95°C for 5 minutes and then quickly placing on ice. As a result of fast cooling, the DNA does not have time to anneal and partially remains in a single-stranded form.

Samples with a volume of 30 μl were prepared containing different concentrations of ATP (0 mM, 2 mM and 10 mM). Each sample contained single-stranded or double-stranded DNA at a concentration of 100 ng/ml and RecN at the rate of 10^3^ molecules per 1 DNA molecule. Samples that did not contain RecN served as controls.

Electrophoresis was then carried out in a 0.7% agarose gel based on Tris-acetate buffer in a standard manner. After electrophoresis, the gel was incubated in TAE containing the intercalating dye ethidium bromide at a concentration of 0.01% to visualize the DNA.

## 3. Results and discussion

### 3.1 RecN increases the frequency of recombination exchanges in *E. coli* by approximately 3 times

We assessed the influence of the RecN on the efficiency of the recombination process in cells, which was determined by the yield of recombinants after conjugation for 30 minutes of the donor KL227 with the recipient AB1157 pT7 recN, carrying plasmids with the *recN* gene. The yield of Thr+ recombinants was normalized to the number of donor cells in the conjugation. The frequency of recombination exchanges can be expressed through the average statistical distance between two neighboring recombination exchanges, which is calculated using the modified Haldane formula, which includes the linkage between the Thr+ selectable and non-selectable Leu+ markers obtained during the experiment.

As a result, it was found that the RecN protein, when induced *in vivo*, increases the frequency of recombination exchanges by approximately 3 times compared to the control, which indicates its increased recombination activity.

### 3.2 RecN demonstrate DNA-dependent ATPase activity

The detailed characterization of *E. coli* RecN *in vitro* properties, which was possible thanks to the successful isolation of highly purified RecN and RecN::mCherry at high concentrations, was carried out. The need to study the *in vitro* properties of *E. coli* RecN is caused, in particular, by the lack of quantitative data on the ATPase activity, however qualitative assessments have been carried out [9]. At the same time, analysis of the bacterial RecN protein sequence shows the presence of two well-known motifs (Walker A and Walker B) involved in ATP binding and hydrolysis. Also, these motives have significant homology to SMC family proteins. To assess the ability of the RecN protein to hydrolyze ATP in real time, we used a spectrophotometric method described above.

As can be seen from Figure 1, *E. coli* RecN has both intrinsic and single-stranded DNA-dependent ATPase activity. Based on linear regression of the time dependencies, the rate of ATP hydrolysis catalyzed by the RecN in the absence and presence of poly(dT) was determined. The corresponding values of the catalytic constant, calculated as the ratio of the observed rate of ATP hydrolysis to the concentration of the RecN protein in the reaction, turned out to be equal 0.13 ± 0.01 min-1 (N = 3) and 0.24 ± 0.02 min-1 (N = 3) in the absence and presence of poly(dT), respectively, and close to constant estimates in [9].

**Fig. 1.**
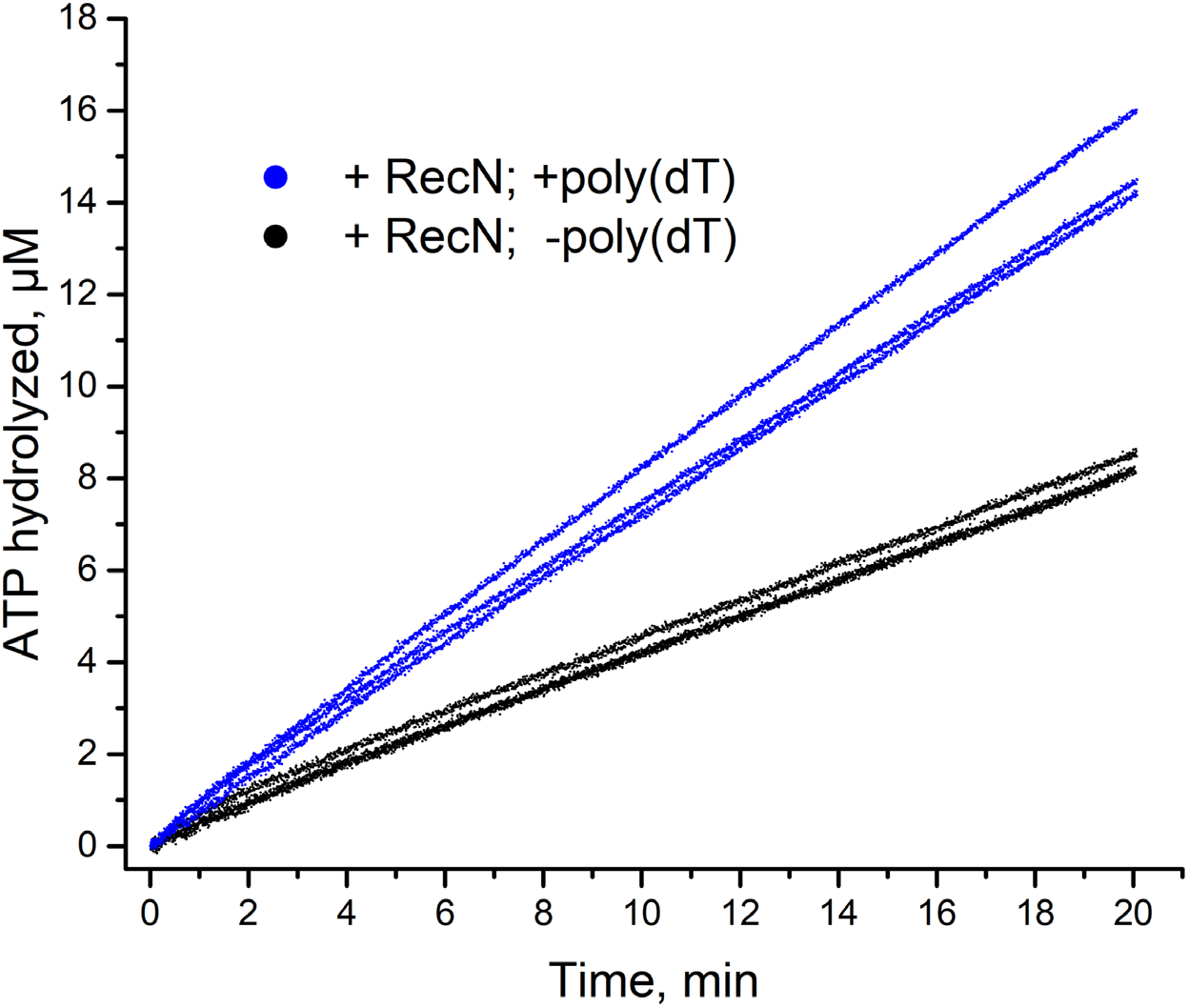
Measurement of *E. coli* RecN ATPase activity in the absence and presence of ssDNA. The formation of ADP during ATP hydrolysis was chemically coupled with the oxidation of NADH to NAD+ and was detected by a decrease in absorption at a wavelength of 380 nm

### 3.3 Under conditions of RecN excess, gel shift data shows its binding ability to both dsDNA and ssDNA

The DNA-binding ability of the RecN was also evaluated using agarose gel electrophoresis of DNA-protein complexes. The change in the electrophoretic mobility of DNA upon binding of RecN to ssDNA and dsDNA was assessed by the smearing of DNA bands and the shift of these bands to higher molecular weight area. Figure 2 shows an ethidium bromide-stained agarose gel loaded with circular ssDNA (samples I-IV) and circular dsDNA (samples V-VIII) in presence and absence of RecN and ATP. It could be concluded that in excess of RecN (10^3^ molecules per 1 DNA molecule) it binds to both dsDNA and ssDNA. Based on band blur efficiency it can be concluded that RecN binding to ssDNA is more pronounced. Also, binding improves significantly with increase in ATP concentration.

**Fig. 2.**
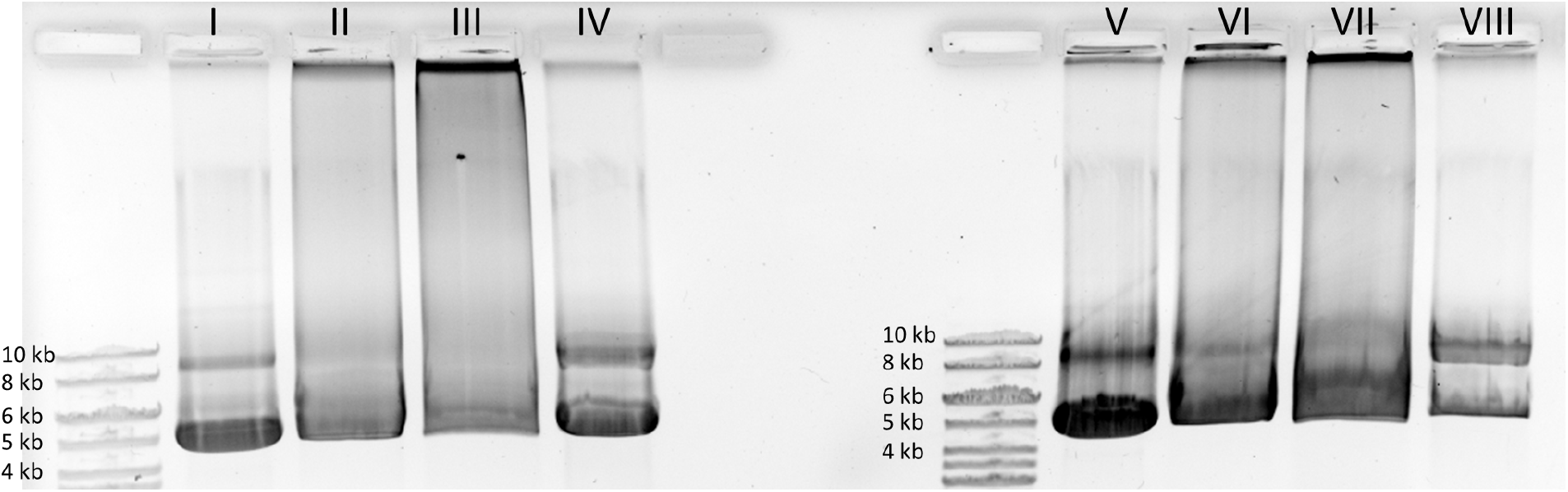
Analysis of DNA electrophoretic mobility in the presence of RecN in a 0.7% agarose gel. I – ssDNA. II – ssDNA, RecN and 2mM ATP. III – ssDNA, RecN and 10mM ATP. IV – ssDNA and RecN. V – dsDNA. VI – dsDNA, RecN and 2mM ATP. VII – dsDNA, RecN and10mM ATP. VIII – dsDNA and RecN

### 3.4 Fluorescence microscopy of single DNA in presence of RecN::mCherry revealed its preferential binding to ssDNA

To obtain more detailed information about the interaction of *E. coli* RecN with DNA, additional experiments were performed using fluorescently labeled RecN::mCherry. Two approaches were used for that. One of them is based on the molecular combining technique described above. Application of this method made it possible to observe, using fluorescence microscopy, protein-DNA complexes formed when RecN::mCherry is incubated with individual DNA molecules straightened on a polystyrene-coated coverslip.

As can be seen from Figure 3a dsDNA molecules stained with the YoYo-1 dye are not associated with the RecN::mCherry, which is located randomly on the glass and did not have pronounced colocalization with dsDNA. In the case of ssDNA, which could not be stained with YoYo-1, a conclusion about the DNA-protein interaction can be made based on fluorescent images analysis, from which it is clear that RecN::mCherry itself has weak fluorescence (Figure 3c, RecN::mCherry), and with the selected contrast is almost invisible. But if RecN::mCherry is added to ssDNA, presumably ssDNA-RecN complexes are formed (Figure 3b, RecN::Cherry + ssDNA). At the same time, in the control ssDNA sample without the protein (Figure 3c, ssDNA), no fluorescence is observed.

**Fig. 3.**
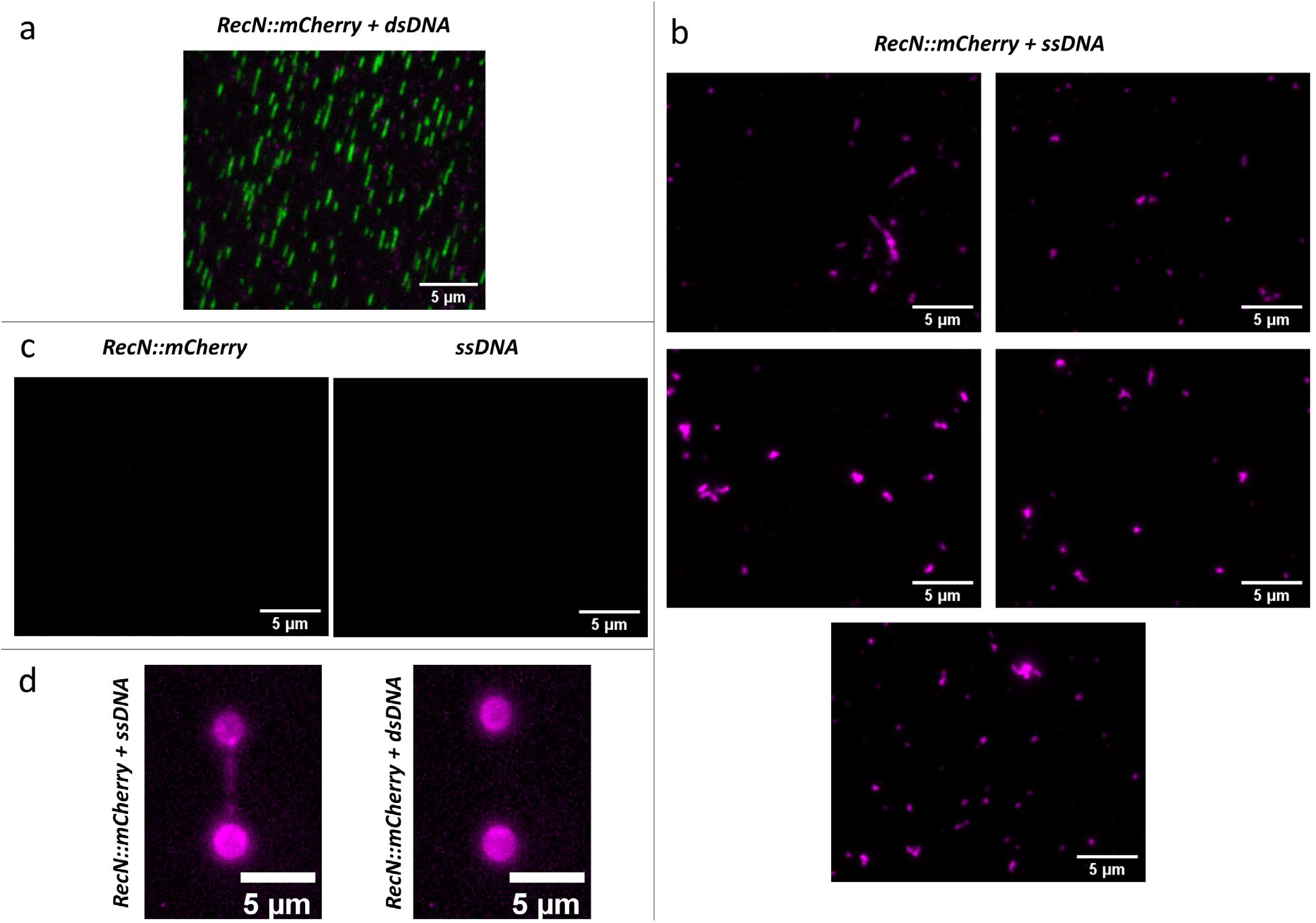
Fluorescence images of single DNA molecules straightened on a glass surface: (a) RecN::Cherry + dsDNA. (b) RecN::Cherry + ssDNA. (c) control – RecN::Cherry on a substrate and ssDNA on a substrate. All a-c fluorescence images have the same contrast. Green – DNA stained with intercalating YoYo-1 dye, magenta – RecN::mCherry. (d) Fluorescence images of single ssDNA and dsDNA molecules tethered between two microspheres using optical traps after incubation with RecN::mCherry

However, in such experiments, DNA and RecN::mCherry are incubated together beforehand molecular combing, so RecN binding could potentially influence the efficiency of anchoring of DNA on glass. The gel shift data (Figure 2) show that a part of the RecN-DNA does not enter the gel, i.e. there could be large complexes. It is possible that such complexes do not bind to the glass during molecular combing.

To clarify this point and to get additional confirmation that RecN::mCherry interacts mostly with ssDNA optical trapping experiments were performed. It was shown that in the case of ssDNA (Figure 3d, RecN::Cherry + ssDNA), the fluorescent signal along the DNA is visible, which corresponds to the formation of a stable RecN::mCherry complex with ssDNA. In the case of dsDNA (Figure 3d, RecN::Cherry + dsDNA), the fluorescent signal along the DNA was at the background level, indicating limited or no binding.

### 3.5 *E. coli* RecN::mCherry localizes at cell poles and in a middle of dividing cell

RecN::mCherry was functional in DNA binding in vitro, we aimed to visualize its localisation in living *E. coli* cells. In the absence of SOS response RecN::mCherry does not colocalize with bacterial DNA. RecN::mCherry has a clear localization pattern and is not distributed evenly throughout the cell. In contrast, it is located predominantly at the cell poles, and also in the area of septum formation in dividing bacteria (Figure 4, RecN::mCherry). Similar results were previously observed using N-terminus GFP-RecN fusion [17,18].

**Fig. 4.**
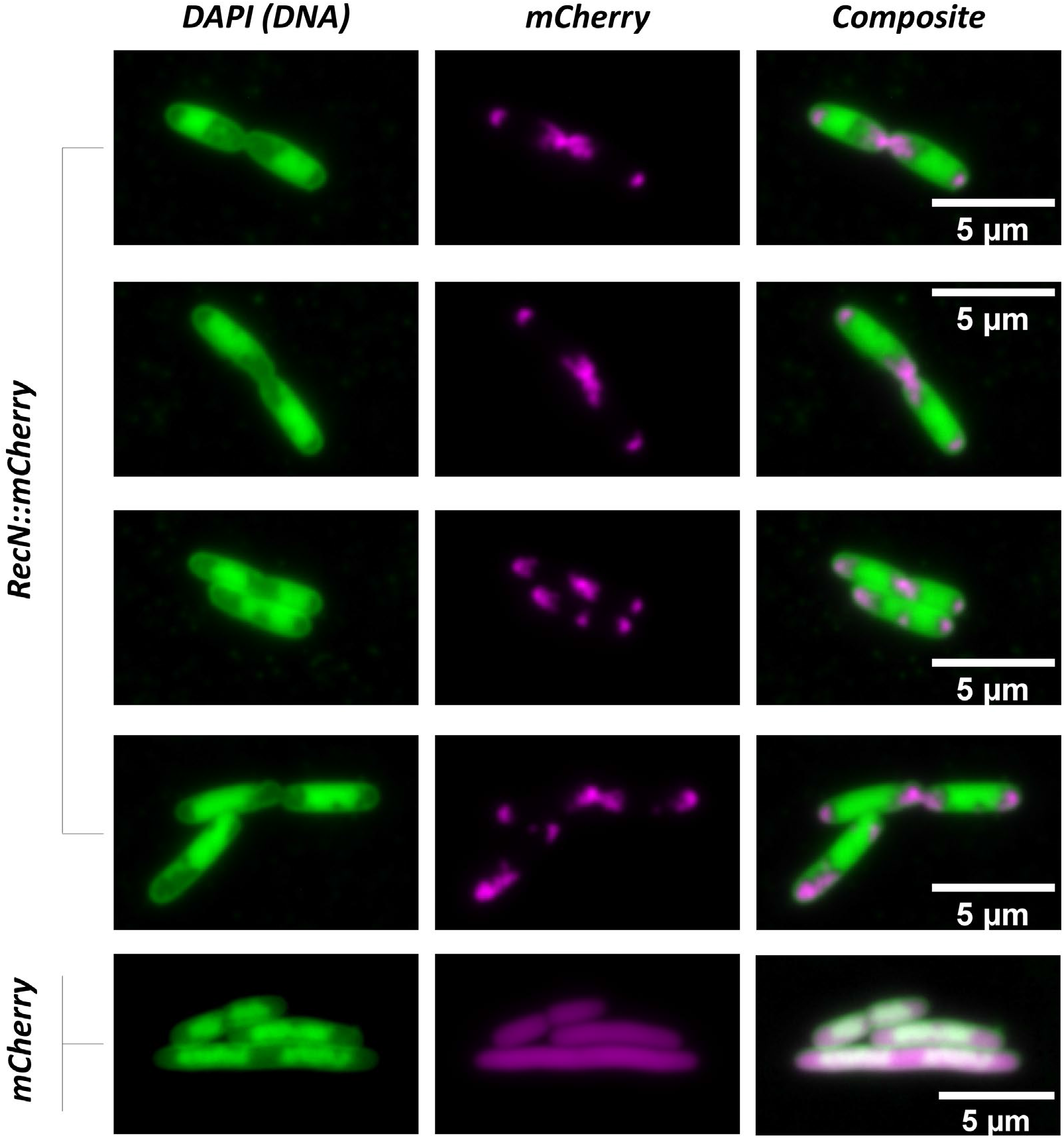
Localization of RecN::mCherry, mCherry and DNA stained with intercalating DAPI dye in living *E. coli* cells. RecN::mCherry and mCherry are shown in magenta, DNA stained with DAPI is shown in green

One can imagine that in the absence of DNA damage, the RecN would be diffusely located throughout the cell what is true for mCherry (Figure 4, mCherry). But it has very specific localization areas and factors that cause RecN to localize in the described manner in the absence of an SOS response, remain unclear. However the fluorescence data are consistent with the lack of binding to dsDNA in optical tweezers and combing experiments.

To conclude, the results of current study demonstrate that expression of RecN increases *E. coli* recombination rate. *E. coli* RecN has intrinsic and ssDNA-dependent ATPase activity and binds preferentially to ssDNA. The binding of RecN to DNA is highly dependent on ATP concentration. Functional *E. coli* RecN::mCherry fusion localizes at cell poles and in a middle of dividing cell and does not colocalize with bacterial DNA out the SOS response.

## Author contributions

**Viktoria D. Roshektaeva:** Formal analysis, Investigation, Writing – original draft. **Aleksandr A. Alekseev:** Formal analysis, Investigation, Writing – original draft. **Alexey D. Vedyaykin:** Methodology, Writing – review & editing, Conceptualization. **Viktor A. Vinnik:** Investigation. **Dmitrii M. Baitin:** Methodology, Investigation, Conceptualization. **Irina V. Bakhlanova:** Methodology, Investigation, Writing – original draft. **Georgii E. Pobegalov:** Methodology, Conceptualization, Writing – review & editing. **Mikhail A. Khodorkovskii:** Methodology, Conceptualization, Writing – review & editing. **Natalia E. Morozova:** Funding acquisition, Methodology, Conceptualization, Investigation, Formal analysis, Data curation, Writing – review & editing, Writing – original draft.

## Acknowledgements

The research was supported by Russian Science Foundation, project No. 22-74-00072 to NEM. The work was carried out using scientific equipment of the Center of Shared Usage “The analytical center of nano- and biotechnologies of SPbPU”.

